# Minocycline enhances antimicrobial activity of unpolarized macrophages against *Acinetobacter baumannii* while reducing the inflammatory response

**DOI:** 10.1101/2024.09.27.615437

**Authors:** Alberto Daniel Guerra, Cecilia F. Volk, Alex Peterson-Weber, Jason M. Peters, George Sakoulas, John-Demian Sauer, Warren E. Rose

**Affiliations:** Pharmacy Practice & Translational Research Division, School of Pharmacy, University of Wisconsin-Madison, Madison, WI USA; Pharmaceutical Sciences Division, School of Pharmacy, University of Wisconsin-Madison, Madison, Wisconsin, USA; Department of Medical Microbiology and Immunology, University of Wisconsin-Madison, Madison, Wisconsin; Department of Bacteriology, University of Wisconsin-Madison, Madison, Wisconsin, USA; Center for Genomic Science Innovation, University of Wisconsin-Madison, Madison, Wisconsin, USA; DOE Great Lakes Bioenergy Research Center, University of Wisconsin-Madison, Madison, Wisconsin, USA; Division of Host-Microbe Systems & Therapeutics, Center for Immunity, Infection & Inflammation, University of California-San Diego School of Medicine, La Jolla, CA, USA

**Keywords:** tetracycline, macrophage, *Acinetobacter baumannii*, antimicrobial, anti-inflammatory, immune response

## Abstract

Minocycline activity against *Acinetobacter baumannii* (*AB*) *in vivo* is underestimated by standard methods of susceptibility testing. We examined pharmacologic effects of minocycline on primary immunity that may be contributing to the *in vivo* vs. *in vitro* discrepancy of minocycline activity against AB. Minocycline MICs against 10 *AB* strains were compared in standard bacteriologic media (Mueller-Hinton broth, MHB) and physiologic (RPMI) media. Macrophages were pretreated with minocycline or comparator antibiotics before *AB* co-culture. Macrophage cytokine production and phagocytosis of *AB* were measured without and with pre-treatment with minocycline. Two to eight-fold reduction in minocycline MIC against 10 *AB* strains occurred in RPMI compared to MHB, which was more pronounced than other antibiotic classes. Macrophages pretreated with 1, 5, 10, 30, 50, and 100 μg/mL minocycline before bacterial co-cultures significantly decreased *AB* inoculum at 6 hours of co-culture in a dose-dependent manner, with no bacterial colonies observed from co-cultures with macrophages pretreated with 30 μg/mL or more of minocycline. Macrophages pretreated with minocycline for 24 hours before zymosan stimulation led to significantly higher levels of phagocytosis. Macrophages treated with minocycline for 24 hours significantly decreased production of IL-6, TNF-α, and MCP-1 in a dose dependent manner. The minocycline *in vivo* efficacy may be attributed to enhanced activity in nutrient-limited, physiologic medium combined with increased macrophage phagocyte efficiency. Incorporating novel assays that recapitulate the *in vivo* environment will be important for understanding the host-pathogen-antibiotic relationship toward a goal of improved future drug discovery and overall treatment strategies against *AB* and other drug-resistant pathogens.

## INTRODUCTION

*Acinetobacter baumannii* (*AB*) is an opportunistic pathogen that portends high patient morbidity and mortality in nosocomial infections, attributed in part to multi-drug resistance (MDR) (1, 2). MDR-*AB* strains are particularly problematic in the intensive care unit and in severe wounds (3-5). Extensively drug-resistant (XDR) AB, including <u>c</u>arbapenem-resistant *<u>A</u>*. *<u>b</u>aumannii* (CR*AB*), brings even greater therapeutic challenges (6). In guidelines on Antimicrobial Resistant Gram-Negative Infections and subsequent updates from the Infectious Diseases Society of America, combination antimicrobial treatment is recommended for CR*AB*. Several agents may be considered in the combination therapy, such as high-dose ampicillin-sulbactam polymyxin B, minocycline, tigecycline, or the newly developed siderophore cephalosporin, cefiderocol (7).

Minocycline is a tetracycline antibiotic that inhibits bacterial protein synthesis through binding to the 30S subunit of the bacterial ribosome. Minocycline use for treatment of CR*AB* is increasing in tandem with the incidence of multi-drug resistant organisms (8). Minocycline activity against *AB* is notable, particularly in its ability to evade the most common tetracycline resistance mechanism (*tetA*) mediated by efflux (9). Clinical studies note positive outcomes using minocycline either monotherapy or as part of combination therapy regimens (10, 11), but the host-response mechanisms involved in these outcomes are poorly understood considering modest minocycline activity *in vitro* against CR*AB*.

While the immune modulating effects of tetracyclines including minocycline are known (12), their relevance in the context of treating MDR-*AB* infections is essentially unexplored. We along with others demonstrated that minocycline improves host immune cell antimicrobial activity and modulates the inflammatory response of monocytes and macrophages (13-16). Additionally, our previous studies have demonstrated that minocycline alters the phenotype of bone marrow-derived mesenchymal stromal/stem cells (MSCs) leading to enhanced wound healing and antibacterial activity against methicillin-resistant *Staphylococcus aureus*, suggesting a non-specific effect on host immunity.(14), implying that this effect may not be pathogen-specific. We posit that minocycline may modulate macrophage antibacterial and inflammatory response against *AB*. In this study, we demonstrate minocycline effect on the unpolarized macrophage inflammatory profile, ability for phagocytosis, and enhanced killing of CRAB.

## RESULTS

### Enhanced minocycline activity against AB in physiologic cell culture media as compared to bacteriologic media

Minocycline appears to be an effective treatment for *AB* out of proportion to its *in vitro* susceptibility and activity (17, 18). This may be due to failure of bacterial growth media typically used to evaluate antibiotic minimum inhibitory concentrations (MICs) to recapitulate the *in vivo* environment. To test if minocycline MICs are affected by medium, ten *AB* strains were evaluated for minocycline, colistin, and meropenem susceptibility in traditional bacterial growth media MHB (Mueller Hinton Broth) versus modified mammalian cell culture medium RPMI + 1% LB (Roswell Park Memorial Institute Medium +1% Lysogeny Broth). As shown in **Table 1**, enhanced minocycline susceptibility in RPMI+1%LB was detected for all strains, leading to a 2-8 fold reduction in MICs compared to MHB. Notably for the CR*AB* strain AB5075, minocycline MIC was 1 µg/ml in MHB and 0.25 µg/ml in RMPI+1%LB. For the comparator antibiotics colistin and meropenem, moderate to no susceptibility changes were found (≤ 2 fold reduced MIC). It is notable that *AB* strains demonstrating intermediate susceptibility or resistance to minocycline in MHB (8 or ≥16 µg/ml per CLSI (19), respectively) were susceptibility (≤4 µg/ml) in RPMI. Thus, it appears that standard bacteriologic media underestimates the potential *in vivo* activity of minocycline.

**Table 1.** Antibiotic susceptibility (μg/ml) of *AB* in traditional growth media compared to mammalian cell growth (RPMI+1% LB). *Indicates clinically derived strain.

| *A. baumannii* | <u>Minocycline</u> |  | <u>Colistin</u> |  | <u>Meropenem</u> |  |
| --- | --- | --- | --- | --- | --- | --- |
|  | MHB25 | RPMI | MHB25 | RPMI | MHB25 | RPMI |
| ATCC 19606 | 0.25 | 0.125 | 2 | 2 | 0.25 | 0.125 |
| AB5075 | 1 | 0.25 | 1 | 1 | 32 | 32 |
| AB248* | 0.5 | 0.25 | 0.5 | 0.25 | 32 | 16 |
| AB250* | 0.25 | 0.125 | 128 | 64 | 32 | 16 |
| AB243* | 0.5 | 0.125 | 0.25 | 0.25 | 1 | 1 |
| AB253* | 2 | 0.5 | 8 | 8 | 64 | 32 |
| AB244* | 8 | 1 | 16 | 16 | 32 | 16 |
| AB610* | 8 | 1 | 0.5 | 0.5 | 16 | 16 |
| AB613* | 8 | 0.5 | 0.5 | 0.5 | 16 | 16 |
| AB625* | 8 | 1 | 0.5 | 0.25 | 4 | 4 |
| AB741* | 32 | 4 | 1 | 1 | 128 | 32 |

### Dose-limiting toxicity of minocycline

Macrophage viability with increasing minocycline concentrations (sub-to supra-therapeutic) was determined to establish non-toxic range to use in the macrophage immune mediated studies. Unpolarized THP-1 macrophages remained viable when treated with up to 100 μg/mL minocycline compared to untreated macrophages after 24 hours of culture (Supplemental **Fig. S1A-B**). There was a higher number of necrotic macrophages when treated with 50 and 100 μg/mL minocycline compared to untreated macrophages after 24 hours of culture (**Fig. S1C**). Using an MTT assay, macrophage metabolic activity was reduced when treated with 100 μg/mL minocycline compared to untreated macrophages and compared to macrophages treated with 50 μg/mL minocycline after 24 hours of culture (**Fig. S1D**). We conclude that minocycline is non-toxic to macrophages up to 50 μg/mL.

### Antimicrobial activity of minocycline-enhanced macrophages against AB

To understand the dose-response relationship between minocycline and macrophage activity against *AB*, we pretreated macrophages with 1, 5, 10, 30, 50, and 100 μg/mL minocycline for 24 hours before bacterial co-culture. we pretreated THP-1 macrophages with 1, 5, 10, 30, 50, and 100 μg/mL minocycline for 24 hours before bacterial co-culture. We found significantly decreased *AB* burden at 6 h of co-culture in a dose-dependent manner, with no bacterial colonies observed from co-cultures with macrophages pretreated with 30 μg/mL or more of minocycline (**Fig 1A**). With minocycline-pretreated macrophages as low as 1 ug/mL, we observed significantly decreased colony forming abilities of *AB* compared to untreated macrophages. This effect was not observed with non-tetracycline antibiotics such as meropenem, piperacillin/tazobactam and colistin. THP-1 macrophages pretreated with 30 μg/mL tetracycline did not decrease the colony forming ability of *AB* at 6 hours of co-culture (**Fig 1B**). Mouse macrophages pretreated with 30 μg/mL minocycline also significantly decreased the colony forming ability of *AB* after 6 hours of co-culture demonstrating that the effects of minocycline on macrophage function are not species specific (Supplemental **Fig. S2**). We next evaluated the antimicrobial properties of the minocycline pretreated human macrophage supernatant extracts against *AB*. Supernatant without macrophages significantly decreased *AB* CFU at 6 hours, with no growth observed in supernatant extracts from macrophages pretreated with 5 μg/mL or more minocycline (**Fig. 2**).

**Figure 1.**
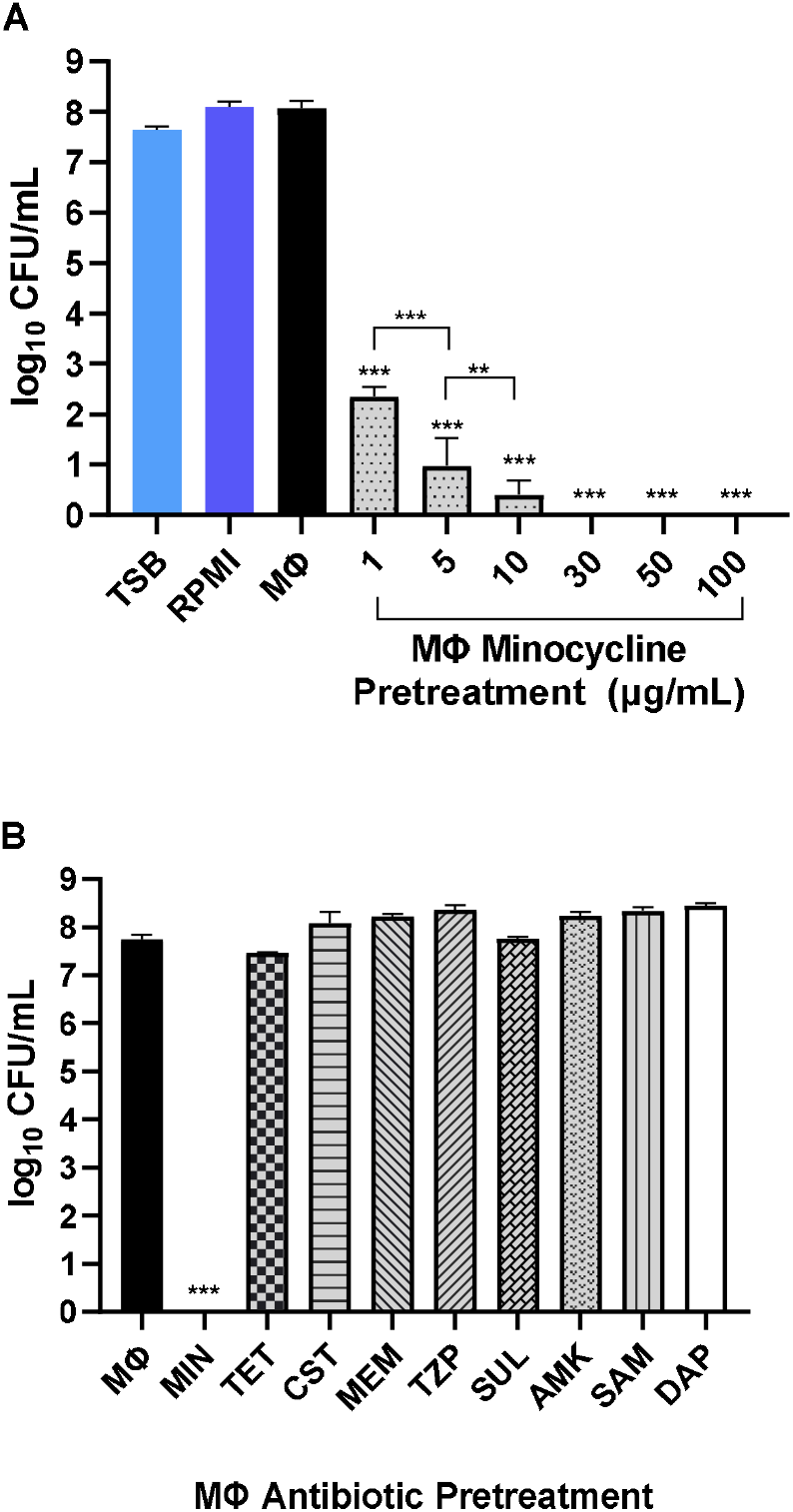
Antimicrobial activity of antibiotic pretreated macrophages against Acinetobacter baumannii at 6 hours of co-culture. A. Colony forming units of *AB* against macrophage co-culture with increasing concentrations of minocycline pretreatment. B. Colony forming units of *AB* against macrophage co-culture after pretreatment across antibiotics classes with concentrations described in materials and methods; TSB, media control; RPMI, media control, MΦ, macrophages in RPMI without drug; Comparisons relevant to MΦ unless otherwise noted; **P<0.01, ***P<0.001.

**Figure 2.**
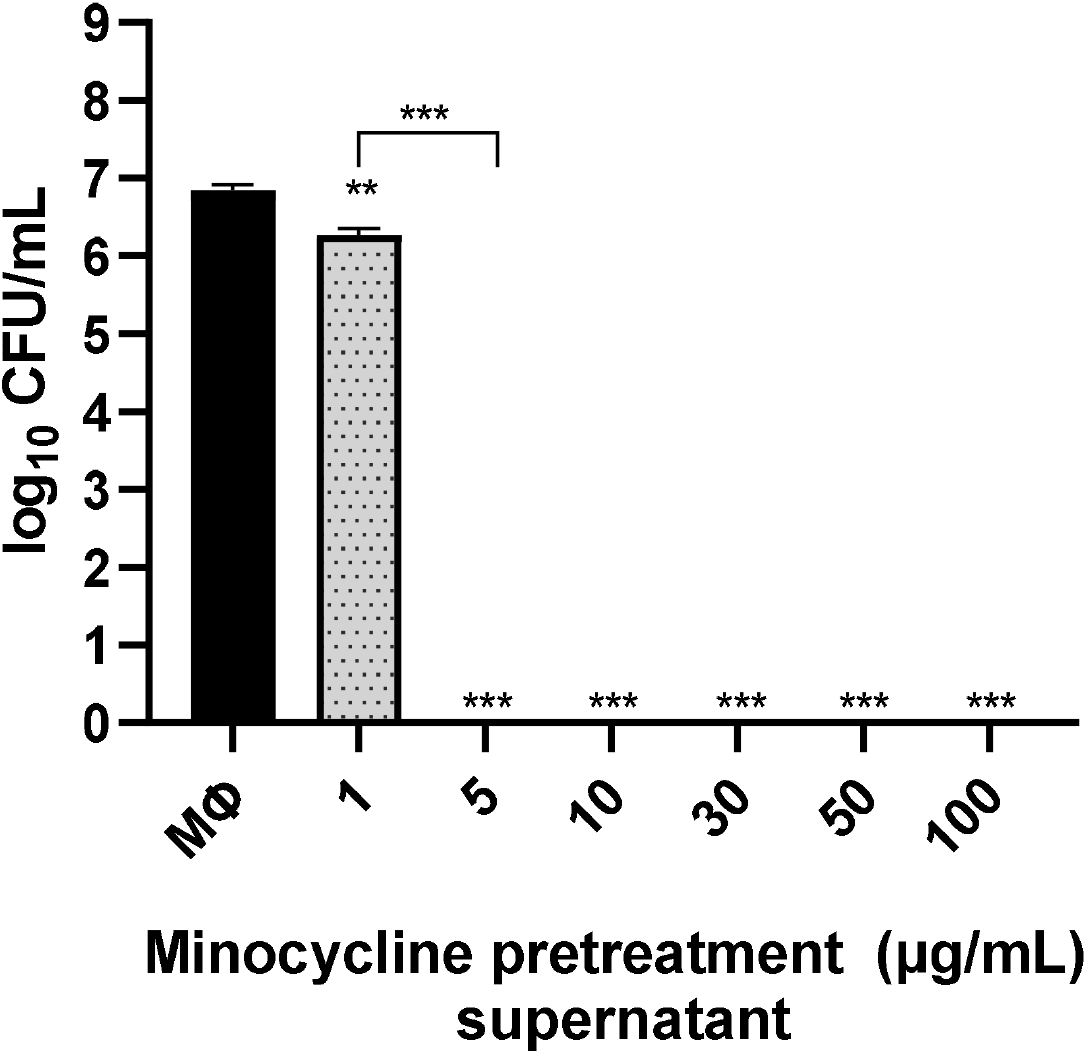
Antimicrobial activity of supernatants from minocycline-treated macrophages against Acinetobacter baumannii at 6 hours. MΦ, macrophage supernatant without drug; Comparisons relevant to MΦ unless otherwise noted; **P<0.01, ***P<0.001.

### Anti-inflammatory properties of minocycline-enhanced macrophages

We sought to investigate if treatment of macrophages with minocycline altered the production of inflammatory factors. Macrophages treated with minocycline for 24 hours significantly decreased baseline production of IL-6, TNF-α, and MCP-1 in a dose dependent manner. Macrophage IL-8 and IP-10 production was significantly lower in macrophages treated with 100 μg/mL minocycline, and 50 and 100 μg/mL minocycline respectively (**Fig 3A**). With AB co-culture stimulation, macrophages pretreated with minocycline for 24 hours had a significant decrease in IL-6, TNF-α, IL-1β, MCP-1, and IP-10 after 6 hours in a dose dependent manner (**Fig. 3B**). Macrophage IL-8 production was significantly lower in macrophages pretreated with 10 μg/mL or more minocycline for 24 hours after 6 hours of *AB* co-culture (**Fig. 3B**). Macrophages pretreated with minocycline for 24 hours demonstrated a significant decrease in nitric oxide production after 6 hours of *AB* co-culture, but there was no difference in the absence of *AB* (Supplemental **Fig. S3**). In assessing anti-inflammatory effects of other antibiotics, macrophages treated with daptomycin or tetracycline for 24 hours also had significantly reduced production of IL-6 and TNF-α (Supplemental **Fig. S4A**). However, in *AB* – macrophage co-cultures, only TNF-α production was reduced with tetracycline (Supplemental **Fig. S3B**). We conclude that inflammatory factors are broadly reduced by minocycline treatment of macrophages even in the presence of AB co-culture as an inflammatory stimulus.

**Figure 3.**
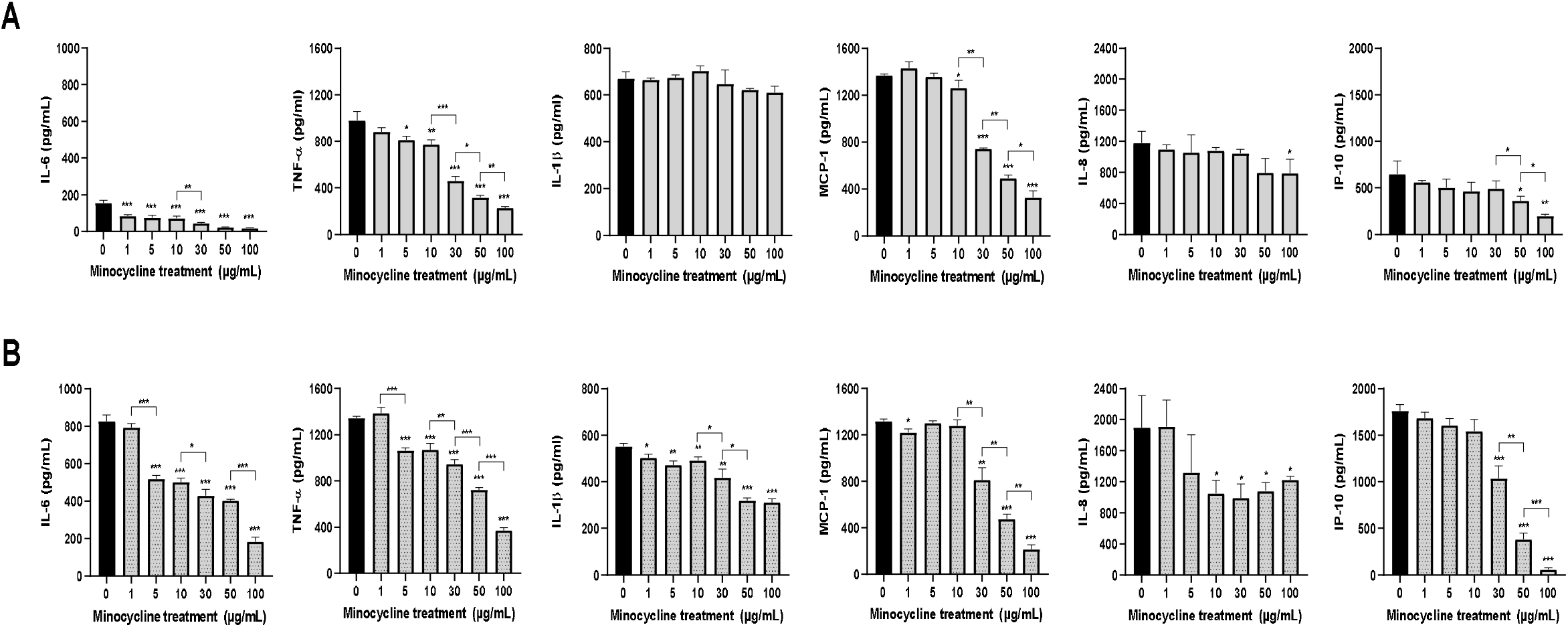
Anti-inflammatory effects of minocycline on unpolarized macrophages. **A**) Production of pro-inflammatory cytokines and chemokines after 24 hours of minocycline treatment; (**B**) Production of pro-inflammatory cytokines and chemokines after 6 hours of co-culture with *AB* following 24 hours of minocycline pretreatment. Comparisons relevant to 0 µg/mL unless otherwise noted; P<0.05, **P<0.01, ***P<0.001.

### Minocycline enhances the phagocytic potential of macrophages

Next, we evaluated the ability of minocycline to enhance the phagocytosis capacity of macrophages. Macrophages pretreated with 30, 50, 100 μg/mL minocycline for 24 hours before zymosan stimulation led to significantly higher levels of phagocytosis of zymosan after 15, 30, 60, and 120 minutes of exposure. Macrophages pretreated with 1, 5, or 10 μg/mL minocycline for 24 hours before zymosan stimulation led to significantly higher levels of phagocytosis of zymosan after 30, 60, and 120 minutes of exposure (**Fig. 4A**). Macrophages pretreated with 10 μg/mL or more minocycline for 24 hours before bacterial co-culture demonstrated significantly higher levels of Beclin1 production after 6 hours of *AB* co-culture (**Fig. 4B**). We conclude that minocycline treatment of macrophages enhances phagocytosis.

**Figure 4.**
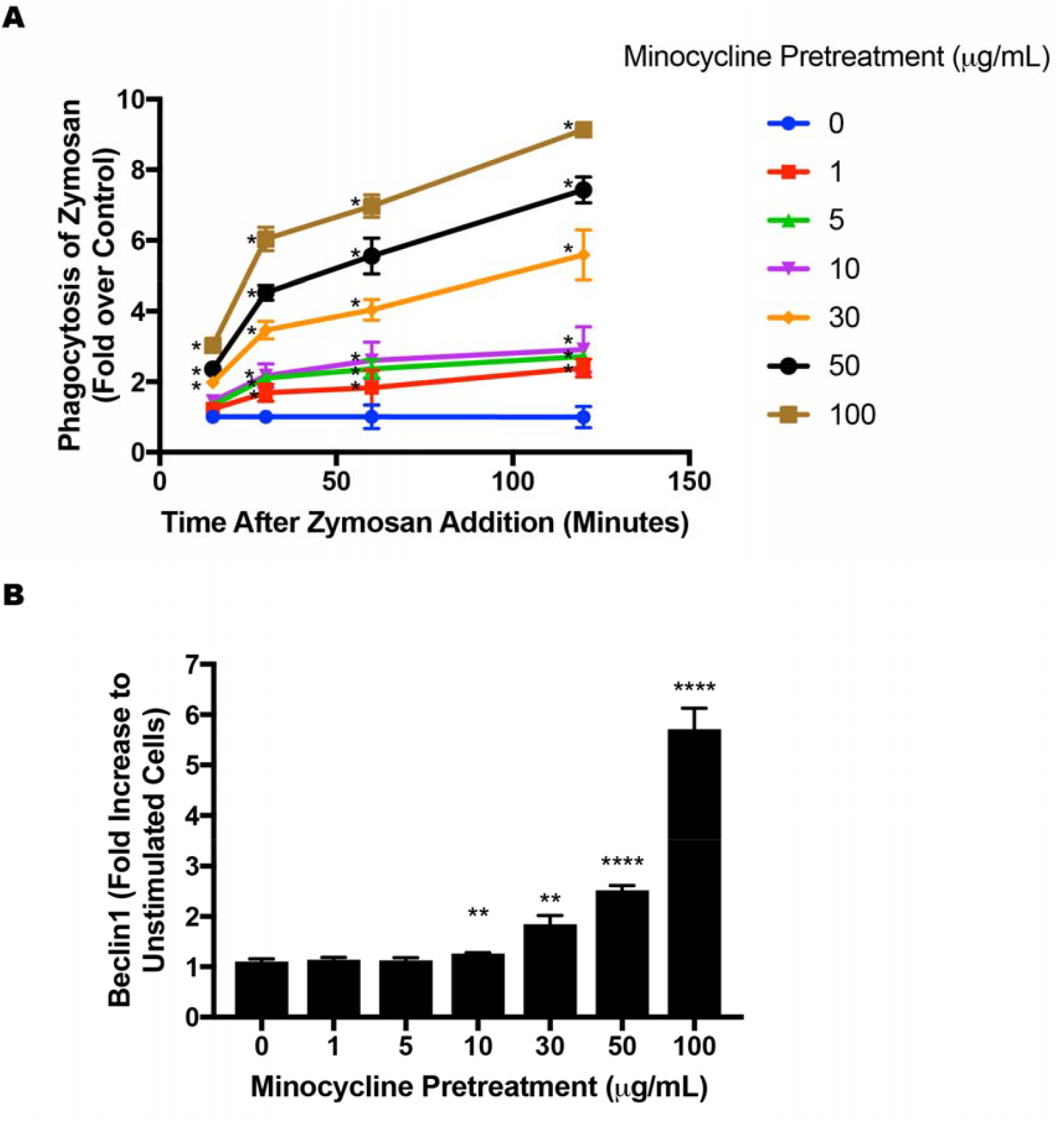
Induction of phagocytosis in unpolarized macrophages by minocycline treatment. **A**) Phagocytosis of zymosan 15, 30, 60, and 120 minutes after 24 hours of minocycline pretreatment. (**B**) Production of beclin1 in cell lysates after 6 hours of minocycline pretreatment followed by 6 hours of co-culture with AB. Comparisons relevant to 0 µg/mL; *P<0.05, **P<0.01, ****P<0.0001.

## DISCUSSION

When susceptibility allows, carbapenems are the most effective treatment option and therefore preferred for *AB* infections (20). However, due to rising carbapenem resistance among *AB*, sulbactam, a β-lactamase inhibitor marketed in combination with ampicillin, has been increasingly relied upon to treat *AB* infections (20). Indeed, sulbactam is a key component of the newly approved antibiotic combination sulbactam-durlobactam to treat CR*AB* (21).

While minocycline is a treatment option for AB, including CR*AB*, its modest bacteriostatic activity and low concentrations in blood raise concerns when used in serious systemic infections (22). However, these concerns are alleviated when outcomes of using minocycline for serious *AB* infections are examined, even in critically ill patients with multiple comorbid conditions and compromised immune systems who respond to treatment with minocycline (23-26). These effects may be attributed to i) evasion of many *AB* resistance mechanisms (9, 25), ii) favorable tissue site pharmacokinetics due to its large volume of distribution (22, 27), and low potential for adverse effects (25) along with non-established off target pharmacologic modification of the host immune response. In this study, we demonstrate that the clinical efficacy of minocycline against AB may also be attributed to the antimicrobial enhancement and inflammatory response reduction of innate immunity.

Minocycline reduces the inflammatory profile in lipopolysaccharide-stimulated THP-1 monocytic cells and macrophages by reducing the production of pro-inflammatory cytokines and the mRNA expression of pro-inflammatory markers (15, 16). These results led us to suspect that additional host environmental factors present *in vivo* during therapy augment minocycline activity against *AB* that are completely missed in routine susceptibility testing in bacteriologic media.

Minocycline is a tetracycline, a dynamic class of molecules that have established structure-activity-relationships as antibacterials, antifungals, antivirals, antineoplastics, as well as functional eukaryotic cell enhancers (13, 14, 28-30). Minocycline differs from tetracycline by the addition of a dimethylamine group on C7 and the loss of a methyl and alcohol group on C6. Despite these very minor differences in chemical structure, we noted a drastic change in potential for macrophage enhancement between the two compounds (30, 31). These small changes in the chemical structure may be important in how minocycline binds and activates its target receptor(s) on macrophages that leads to antimicrobial enhancement. Elucidation the target receptor and downstream cell signaling pathways will be the goal of future studies that could lead to the ability to utilize the activation of these receptors and pathways for alternative therapeutic approaches in multidrug-resistant microbial infections.

Macrophages treated with as low as 1 μg/mL minocycline before removal of minocycline and *AB* co-culture reduced *AB* growth over 6 hours, with the antimicrobial effect increasing with greater concentrations of minocycline. This concentration corresponds to the pharmacokinetic and pharmacodynamics data of patients treated with an IV administration of minocycline for MDR-*AB (10)*. In addition, macrophages treated with minocycline concentrations of 30 µg/mL or higher completely prevented *AB* growth.

The mechanism behind this antimicrobial effect has not been completely elucidated but appears to be multifactorial. Minocycline induces internalization of zymosan in macrophages, suggesting that macrophage exposure to minocycline leads to an increase in macrophage phagocytosis of *AB*. This is supported by the increase in beclin-1 production when macrophages were stimulated by minocycline, as beclin-1 is an autophagy-related gene that is known to be upregulated by minocycline (32). However, we also found antimicrobial effects of supernatant collected after the co-culture of minocycline-treated macrophages with *AB*, suggesting factors such as antimicrobial compounds produced by macrophages after minocycline stimulation beyond phagocytosis are involved.

The anti-inflammatory effects of minocycline warrant discussion as well given what is known about acute lung injury in the pathophysiology of pneumonia. Specifically, clearance of bacterial and/or viral pathogens from the alveolar space may be accompanied by severe inflammation that may be highly successful in pathogen clearance, but unfortunately carries collateral damage of the fragile alveolar membrane which is critical in gas exchange. Therefore, attention is frequently given to immunomodulation in pneumonia as evidenced by literature on the benefit of glucocorticoids in severe bacterial pneumonia (33), the use of glucocorticoids (34) and targeted anti-IL-6 inhibition (35) in treating COVID-19, as well as data showing survival benefit of adding doxycycline to backbone cephalosporin therapy in treating community-acquired bacterial pneumonia (36). This study extends the appreciation of the potential benefit of anti-inflammatory effects of minocycline specifically in the macrophage interaction with *AB*. Of note in this study, minocycline reduced the production of inflammatory cytokines from macrophages in a dose dependent manner with and without co-culture with *AB*. These cytokines (IL-6, TNF-α, IL-1β, MCP-1, IL-8, IP-10) play a significant role in lung inflammation, pathophysiology, and clinical sequalae of pneumonia. It appears that antibiotic MIC against a pathogen is just one of many considerations in selecting optimal antimicrobial therapy for serious infections and overall *in vivo* efficacy is affected by off-target antimicrobial effects on the reduction the host inflammatory response.

In summary, three separate immune-modulating effects of minocycline were observed in this study: i) an increase in macrophage phagocytosis, ii) alterations in inflammatory cytokine production, and iii) macrophage secretion of an antimicrobial compound or compounds. Coupled with the fact that physiologic media that better recapitulates the *in vivo* environment significantly lowers minocycline MIC against *AB*, multiple reasons appear to explain why clinical data of minocycline treating *AB* infections exceed what *in vitro* assays would predict. Our future studies are focused on identifying the antimicrobial compound secreted by macrophages after minocycline exposure. In demonstrating immune-modulating mechanisms of minocycline, this study highlights the limitations of utilizing traditional susceptibility testing methods that do not account for the host immune system. Incorporating novel assays that recapitulate the *in vivo* environment will be important for understanding the host-pathogen-antibiotic relationship toward a goal of improved future drug discovery and overall treatment strategies against *AB* and other drug-resistant pathogens.

## MATERIALS AND METHODS

### Bacterial strain, media, and antibiotic susceptibility

*AB* strain 5075 was used in for *in vitro* experiments of antibiotic and co-culture activity. (2). In addition, control ATCC strain 19606 and nine clinical multi-drug resistant strains were evaluated for antibiotic susceptibility in bacterial growth media (cation adjusted MHB) and modified mammal cell culture media (RMPI+1% LB). *AB* routine growth and strain maintenance were carried out in tryptic soy agar/broth (TSA/TSB) media (Becton Dickinson, Sparks, MD) unless noted otherwise for specific assays. Antibiotics were purchased from Sigma-Aldrich (St. Louis, MO) and reconstituted and diluted according to manufacturer’s instructions. All antibiotics were freshly prepared and used the day of the experiment.

### Toxicity of macrophages exposed to minocycline

Macrophages were plated in 48 well plates as described in detail below and treated with escalating concentration of minocycline at 0, 1, 5, 10, 30, 50, or 100 μg/mL for 24 hours. After minocycline treatment, supernatants were removed for soluble factor analysis and a LIVE/DEAD® (Invitrogen) stain was applied with subsequent imaging and fluorescence quantification followed by a CellTiter-Blue® (Promega, Madison, WI) stain with subsequent quantification as previously described (28).

### Monocyte culture, stimulation into macrophages, and antibiotic treatment

Human acute monocytic leukemia cell line THP-1 were purchased from American Type Culture Collection (ATCC) and maintained according to ATCC guidelines cultured in RPMI 1640 medium with GultaMAX™ supplement and HEPES buffer (ThermoFisher Scientific) at 10% fetal bovine serum. Media was changed every 3-4 days. For stimulation into macrophages, monocytes were plated at 500,000 cells/well in a 48 well tissue culture plate (VWR International, Randor, PA) in media at 320 nM phorbol 12-myristate 13-acetate (PMA, Sigma-Aldrich) for 24 hours (29, 37). After stimulation into macrophages, PMA-containing media was removed and replaced with fresh media as above or media-containing antibiotic (total volume 500 μL). Macrophages were treated with antibiotic for 24 hours before use in co-culture experiments. Minocycline pretreatment was evaluated at concentrations of 1, 5, 10, 30, 50, and 100 μg/mL. Macrophage pretreatment with comparative antibiotics included tetracycline (TET, 30 μg/mL), colistin (CST, 2 μg/mL), meropenem (MEM, 50 μg/mL), piperacillin/tazobactam (TZP, 200 μg/mL), sulbactam (SUL, 85 μg/mL), amikacin (AMK, 30 μg/mL) ampicillin/sulbactam (SAM, 150/85 μg/mL) and daptomycin (DAP, 10 μg/mL) using concentrations representing achievable therapeutic levels in serum or tissue.

To evaluate effects in a non-human macrophage line, mouse bone marrow-derived macrophages were harvested as previously described (38) and cultured in the same media on non-tissue culture polystyrene plates (VWR International). Similarly, two days before co-culture with *AB*, macrophages were removed from plates with ice-cold PBS for 10 minutes and plated at 250,000 cells/well in a 48 well tissue culture plate (VWR) for 24 hours before antibiotic treatment and co-culture experiments as described above.

### Acinetobacter baumannii culture and co-culture with macrophages

*AB* 5075 culture vials stored at -80°C were thawed and grown on Luria Bertani agar overnight. *AB* colonies were then added to a tissue culture glass test tube with 2ml sterile double-distilled water to achieve 0.5 McFarland standard. Next, 1ml of this solution was then added to 29ml of TSB growth medium and incubated at 37°C on a rotator until a spectrophotometer (Nanodrop 2000x UV-vis Spectrophotometer, Thermo Scientific) OD_600_ of 0.1 (∼1×10^5^ CFU/mL) was achieved.

To collect antibiotic-pretreated macrophages, the cell culture treatment media was removed from the macrophage cultures and each well was washed 3x with 500μL of PBS to completely remove antibiotic from the culture wells and retain the pretreated macrophages.

Next, 500μL of fresh RPMI media without antibiotic was added to all wells. *AB* (starting at 1×10^5^ cfu/ml) was then added to macrophage cultures treated with or without antibiotic. At sample time points of 6 and 24 hours, 50μL for co-culture supernatant samples was removed, serially diluted in 0.9% NaCl and plated on TSA for colony enumeration as described previously (28).

Macrophages are capable of secreting innate host defense peptides, so we also evaluated the supernatants of the cell culture for antibiotic-stimulated antimicrobial effects against *AB*. Cell culture media from the pre-treated macrophages above were collected in 1.5 mL microcentrifuge tubes and spun at 12,000x g for 15 minutes. The supernatant was then transferred to a clean microcentrifuge tube. 500 µL of the extracted supernatant was added to 1.5 mL microcentrifuge tubes and AB5075 was added at 1×10^5^ cfu/ml for 6 hours to evaluate bacterial killing.

### Macrophage antibacterial response analyses

Supernatants from i) 24 hour macrophage pretreatment and ii) the 6 hour co-cultures of pretreated macrophages with *AB* were collected as described above and analyzed for production of soluble factors using a DuoSet® ELISA Development System (R&D Systems, Inc., Minneapolis, MN) with corresponding assay antibodies and standards as described previously (13). Nitric oxide levels were determined using Griess Reagent (Sigma Aldrich) as previously described (29).

### Macrophage phagocytosis analysis

Phagocytosis potential of minocycline-treated macrophages was determined using pHrodo™ Red Zymosan Bioparticles™ Conjugate (ThermoFisher Scientific). THP-1 cells were plated at 100,000 cells/well in 96 well tissue culture plates, stimulated into macrophages, and treated with either 0, 1, 5, 10, 30, 50, or 100 μg/mL minocycline for 24 hours as described above. After pretreatment, cells were washed as described above and treated with Zymosan Bioparticles following the manufacturer’s instructions with analysis at λ_excitation_= 560 nm and λ_emission_= 585 nm at 15, 30, 60, and 120 hours after treatment with Zymosan Bioparticles with appropriate controls on Synergy H1 Hybrid Microplate Reader (Biotek Instruments, Winooski, VT, USA).

The levels of the regulator of autophagy Beclin 1 in macrophages were determined by washing cells with PBS after co-culture, lysing cells with RIPA lysis buffer (ThermoFisher Scientific) with Halt™ Protease and Phosphatase Inhibitor Cocktail (ThermoFisher Scientific) with quantification for Beclin1 by ELISA (Lifespan Biosciences, Seattle, WA).

### Statistical analyses

All assays were conducted with at least three replicates per condition. A two-way ANOVA for multiple comparisons was used with a Bonferroni test assessment to determine *p* values for assays. A P<0.05 was considered statistically significant with P<0.05 (*), P<0.01 (**), P<0.005 (***), P<0.001 (****). All analyses were performed with GraphPad Prism 9.0. Error bars represent the standard deviation of the specific group mean.

