## Supplemental Figures for "Minocycline enhances antimicrobial activity of unpolarized macrophages against *Acinetobacter baumannii* while reducing the inflammatory response"

**Supplemental Figure S1.** Supplemental Figure S1. Cytotoxicity of unpolarized macrophages treated with 0 to 100 μg/mL minocycline for 24 hours. (A) Live/Dead images of minocycline-treated macrophages. (B) Quantification of live fluorescence in minocycline-treated macrophages. (C) Quantification of necrotic fluorescence in minocycline-treated macrophages. (D) Quantification of metabolic activity in minocycline-treated macrophages. *P<0.05, **P<0.01, ***P<0.001, ****P<0.0001.

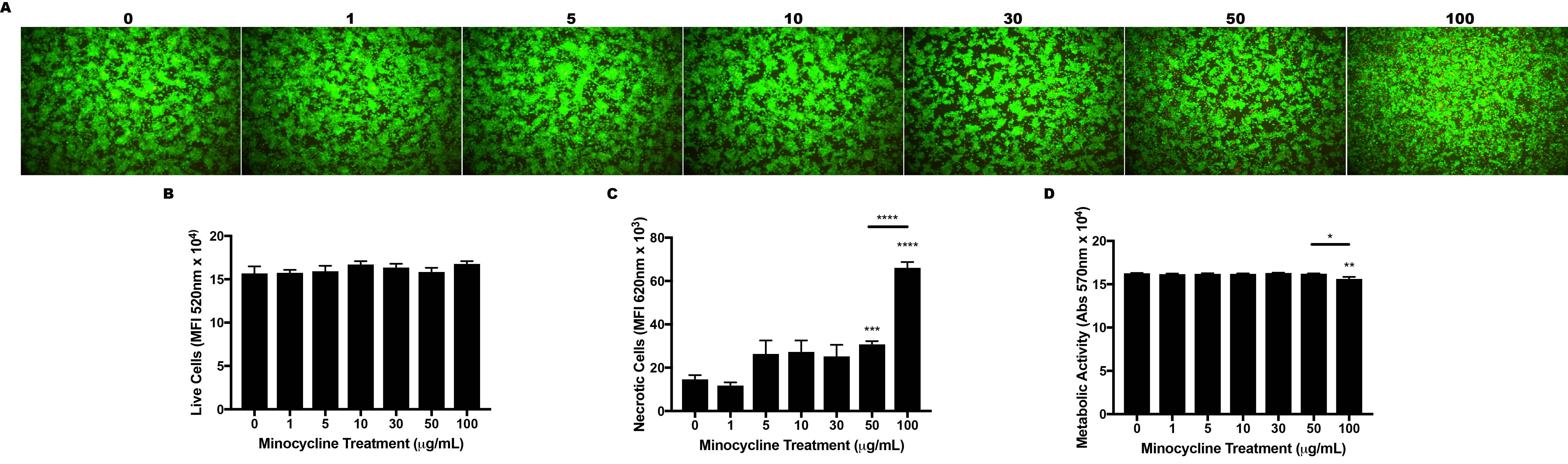

**Supplemental Figure S2.** *AB* growth at 6 hours with pretreatment of murine bone marrow-derived macrophages (M0) with minocycline 30 µg/mL. ***P<0.001

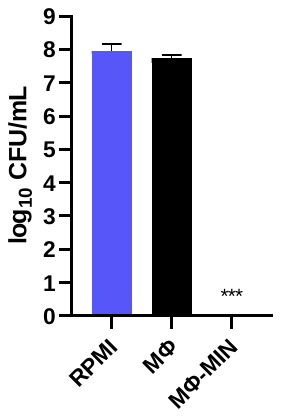

**Supplemental Figure S3.** Nitric oxide production in macrophages pretreated with 30 μg/mL minocycline for 24 hours in the absence of *AB* and with *AB* co-culture. *P<0.05, **P<0.01.

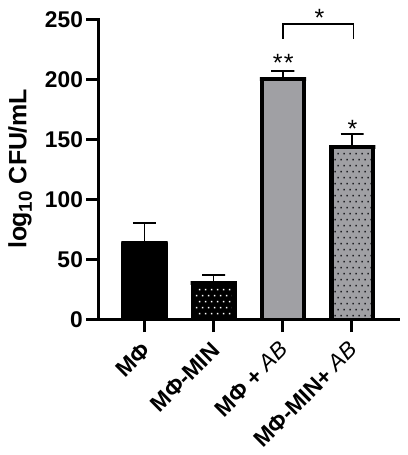

**Supplemental Figure S4.** Anti-inflammatory effects of drug treatments on macrophages. A) IL-6 and TNF-α production from macrophages after 24 hours of drug treatment only B) IL-6 and TNF-α production from macrophages after 24 hours of drug pretreatment in co-culture with AB.

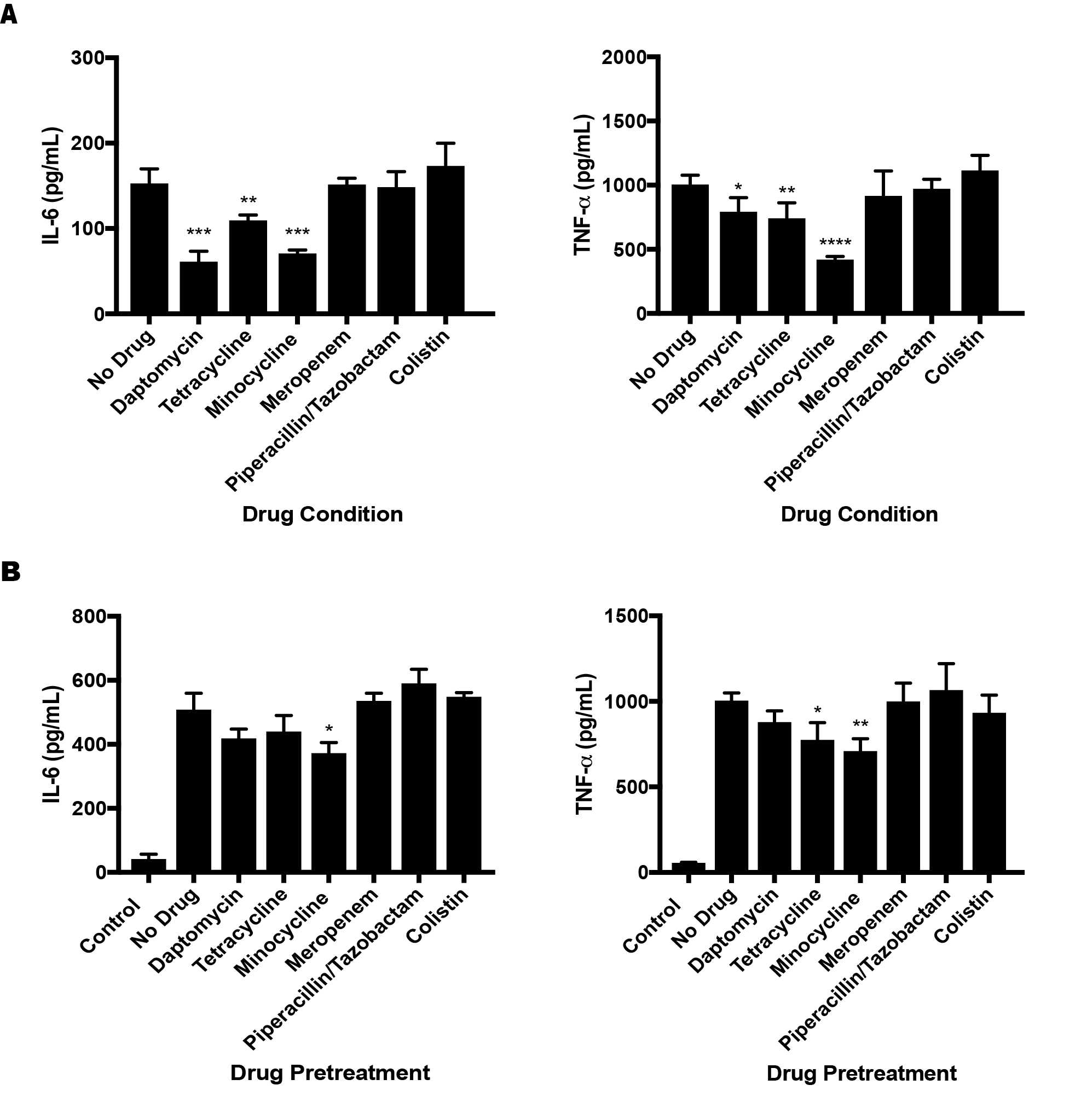
